# Whole genome sequencing-based characterization of mobile genetic elements in *Staphylococcus aureus* isolated from patients in Fort Portal Regional Referral Hospital, Western Uganda

**DOI:** 10.64898/2026.01.13.699411

**Authors:** Moses Luutu Nsubuga, Erick Katagirya, Gavin Ackers Johnson, Kia Praiscillia, Kaddu Arafat, Bwambale Wilberforce, Muwambi William Jonathan, Fahim Yiga, Kalema Leymon, Henry Ssenfuka, Richard Mulondo, Edgar Kigozi, Fred Ashaba Katabazi, Henry M Kajumbula, Chloë E James, David Patrick Kateete

**Author notes:** **Corresponding authors:** (DPK), (MLN).

## Abstract

**Background:** The ability of *Staphylococcus aureus* to evolve through horizontal gene transfer mechanisms aids its success as a versatile pathogen. Mobile genetic elements (MGEs) are linked to potent virulence factors in *S. aureus,* e.g., the Panton-Valentine leukocidin and toxic shock syndrome toxins, as well as antibiotic resistance genes, e.g., *mecA* that encodes methicillin resistance. Despite their clinical relevance, molecular surveillance of MGEs in Africa remains limited. Here, we characterize the MGE repertoire of clinically relevant *S. aureus* isolates from Fort Portal Regional Referral Hospital (FPRRH), western Uganda.

**Methods:** We assembled a total of 40 genome sequences from previously sequenced *S. aureus* isolates cultured from patients (skin wounds, urinary tract, and bloodstream infections) at FPRRH during 2017-2019. spaTyper was used to determine the spa genotypes, while the presence of MGEs was screened and annotated for using PlasmidFinder, PHASTEST, Mobile Element Finder, SCC*mec*Finder, Bakta, MobileOG-db, and IslandViewer tools.

**Results:** Eleven spa types were identified, with spa type t355 predominating. We detected 74 plasmid-derived sequences and 31 insertion sequences. Two SCC*mec* types, SCC*mec* type III and SCC*mec* type IV, were detected, indicating both hospital-associated MRSA (HA-MRSA) and community-associated MRSA (CA-MRSA). Forty-seven intact prophages (all Siphoviridae) were identified, carrying *dfrG, sak*, and *lukPV* genes. A total of 191 genomic islands were detected, and these harbored the virulence, immunoevasion, drug, and heavy metal resistance genes, such as *nuc, tuf, tst, pvl, tet, blaZ,* and *mer* genes.

**Conclusions:** *S. aureus* at FPRRH harbors a diverse and functionally rich MGE repertoire, including genomic islands, prophages, insertion sequences, transposons, and plasmids, that contribute to the dissemination of virulence, AMR, and metal resistance determinants. The coexistence of HA-MRSA and CA-MRSA, as seen in other regions of Uganda, underscores the importance of continued genomic surveillance to inform infection control strategies.

## Introduction

*Staphylococcus aureus* is a leading cause of both community- and healthcare-associated infections worldwide [1]. It is a highly adaptable commensal bacterium capable of causing a broad range of clinical manifestations, from superficial skin infections to severe and potentially fatal diseases such as osteomyelitis, endocarditis, and sepsis [3]. In Sub-Saharan Africa, *S. aureus* infection is particularly concerning as the pathogen ranks as the second most common cause of pneumonia among children [2].

*S. aureus* infections often arise from endogenous reservoirs, with colonization in the anterior nares, skin, and mucosa serving as major sources [1]. Transmission occurs through direct contact or breaches in host barriers, yet the capacity of *S. aureus* to establish fatal infection is strongly influenced by mobile genetic elements (MGEs) such as plasmids, transposons, bacteriophages, insertion sequences, and genomic islands that facilitate horizontal gene transfer and drive acquisition of virulence and antibiotic resistance determinants [7–10]. *S. aureus* pathogenicity islands (SaPIs) encode superantigens such as toxic shock syndrome toxin (TSST) and enterotoxins, while prophages carry Panton–Valentine leukocidin (PVL), enhancing immune evasion and tissue damage through production of hemolysins, Protein A and Aureolysin [6].

The staphylococcal cassette chromosome *mec* (SCC*mec*), a genomic island harboring the *mecA* gene, exemplifies the clinical relevance of MGEs by conferring methicillin resistance and defining methicillin resistance *S. aureus* (MRSA) [11–13]. The strong association between *S. aureus* clonal lineages and distinct MGEs underscores their central role in bacterial evolution, transmission dynamics, and persistence in human populations [10].

According to previous research, *S. aureus* genotypes are associated with specific MGEs [14, 15]. This has necessitated molecular surveillance of the pathogen (i.e., genotyping and characterization of mobile genetic elements). However, due to inadequate infrastructure in many regions of Africa, this surveillance remains restricted [16]. Furthermore, previous research on MGEs is targeted and largely focused on plasmids of resistant strains such as MRSA, leaving many MGEs unexplored. In order to fully understand the evolution and spread of staphylococcal virulence and antibiotic resistance in a low-income setting, we performed whole genome sequencing (WGS) of *S. aureus* clinical isolates at Fort Portal Regional Referral Hospital (FPRRH) in western Uganda and characterized isolates for the mobile genetic elements (MGEs) harbored including the virulence and antibiotic resistance genes encoded.

## Results

### Genomic assembly and annotation

A total of 40 MiSeq-generated paired fastq files were downloaded from the European Nucleotide Archive database, containing raw reads with sizes ranging from 50 to 141 megabytes (MBs). For each *S. aureus* sample, two fastq files, for the forward and reverse reads, were downloaded. FastQC assessment of the raw data indicated 90% of both forward and reverse reads across samples were unique. After adapter trimming and removal of poor-quality reads by fastp, all reads were within the quality score of Q30. The 40 *S. aureus* draft genome sequences were assembled with spades and annotated using PROKKA. The genome sizes after assembly ranged from 2.6 megabase pairs (Mbps) to 4.9 Mbps. Annotation identified at least 1000 coding sequence regions (CDS), 1 tmRNA, 30 tRNA, and 3 rRNA for each genome.

### Staphylococcal protein A (spa) types

Spa typing revealed 11 different spa types across the 40 *S. aureus* genomes. The most prevalent spa type was t355, which was identified in 35% (n=14) of the genomes; 20% (n=8) of the genomes could not be typed and thus were not assigned a spa type (Fig 1).

**Fig 1:**
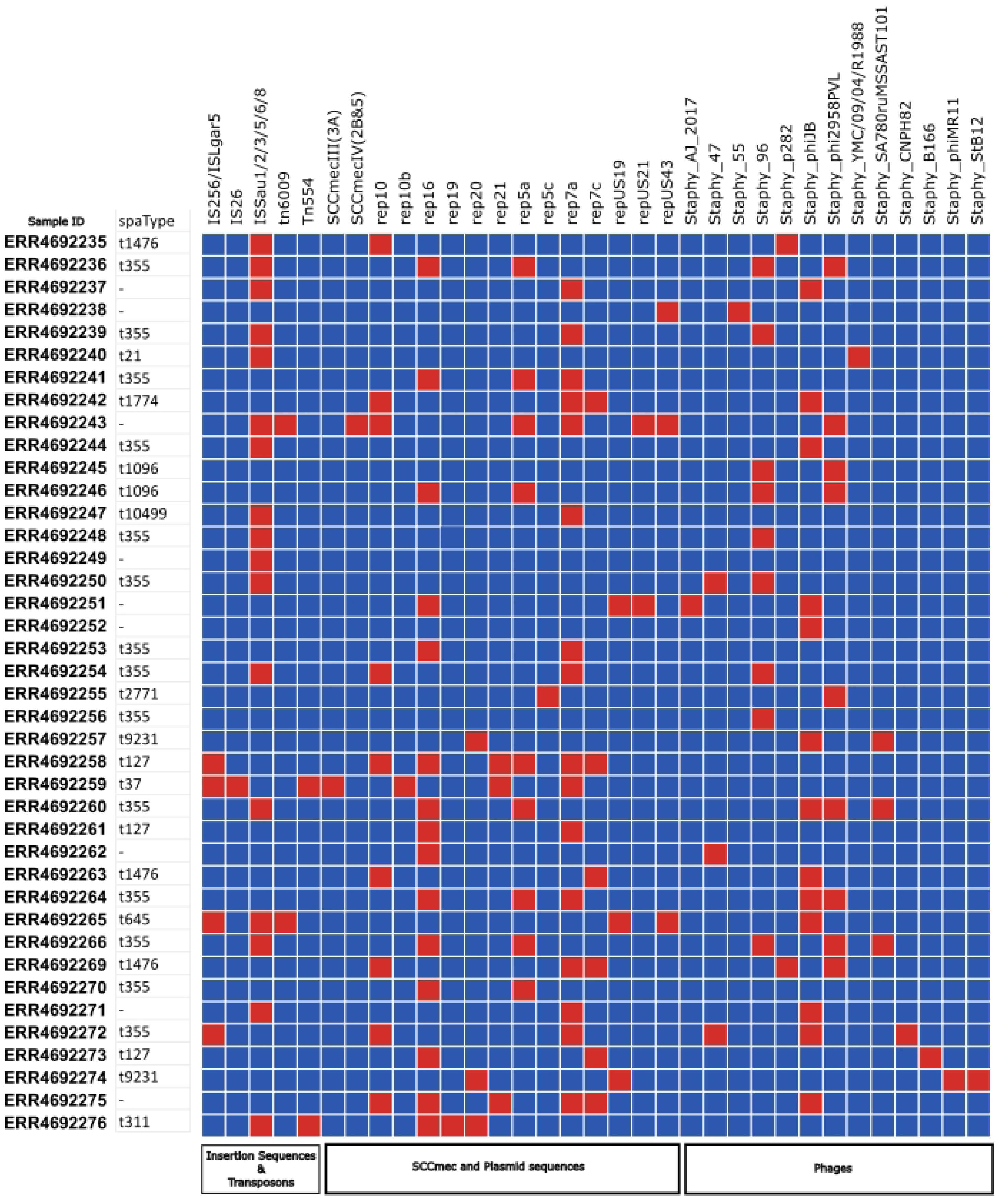
Distribution of spa types and mobile genetic elements (MGEs) identified across 40 assembled Staphylococcus aureus genomes. Red squares indicate the presence, and blue squares indicate the absence of each respective feature in a given isolate.

### MGEs

Plasmid replicons, transposons, SCC*mec*, insertion sequences, and prophages identified are summarized in Fig 1.

#### Plasmids

A total of 74 plasmid-derived sequences were detected across the 40 *S. aureus* genome assemblies, despite the inability to fully reconstruct complete plasmids. Their GC content (25.0%–30.0%) was consistently lower than the chromosomal average. *rep7a* and *rep16* were the most common replicon types, identified in 16 and 15 assemblies, respectively. Three sequences (4%), all *repUS43* plasmids, carried a MOBT family relaxase, indicating mobilization potential. Notably, all plasmid-derived sequences were chromosomally integrated rather than extrachromosomal, frequently adjacent to ORFs encoding conjugal transfer proteins, AMR determinants such as *tet(K)*, and other mobile genetic elements such as transposons and insertion sequences, pointing to historical or ongoing horizontal gene transfer (Figs 2 and 3). The highest plasmid sequence counts occurred in ERR4692243 (n = 7), ERR4692258 (n = 6), and ERR4692275 (n = 5), while 17.5% (7/40) of assemblies contained none.

**Fig 2:**
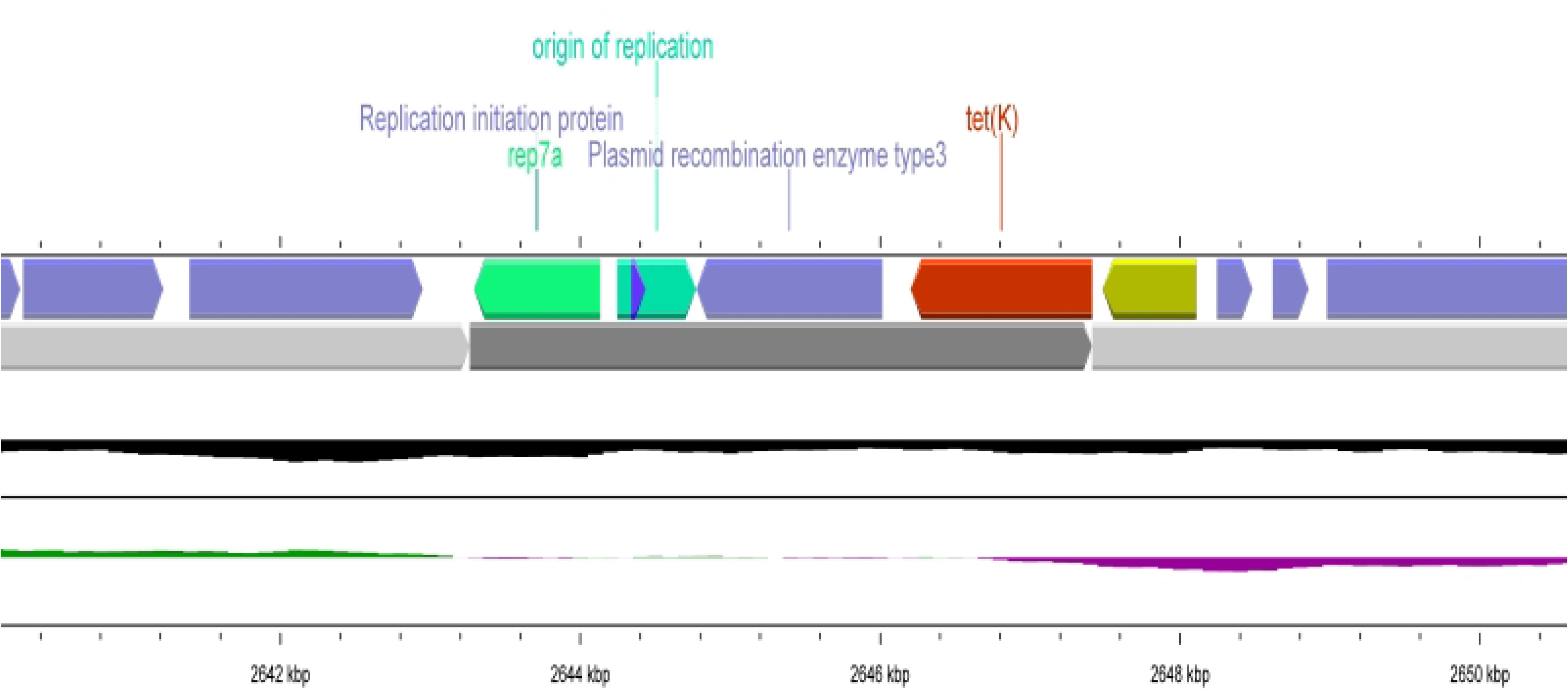
Genomic region of *S. aureus* isolate ERR4692259 illustrating a partial sequence of the plasmid replicon *rep7a* integrated into the chromosome, positioned near an origin of replication (Ori), a type 3 plasmid recombination enzyme, and the tetracycline resistance gene *tet(K)*. Such integration events, often mediated by recombination machinery, highlight potential horizontal gene transfer pathways linking plasmid-chromosome-borne antimicrobial resistance determinants. Tracks from top to bottom: Lane 1 – annotated genes; Lane 2 – contig backbone; Lane 3 – GC content; Lane 4 – GC skew.

**Fig 3:**
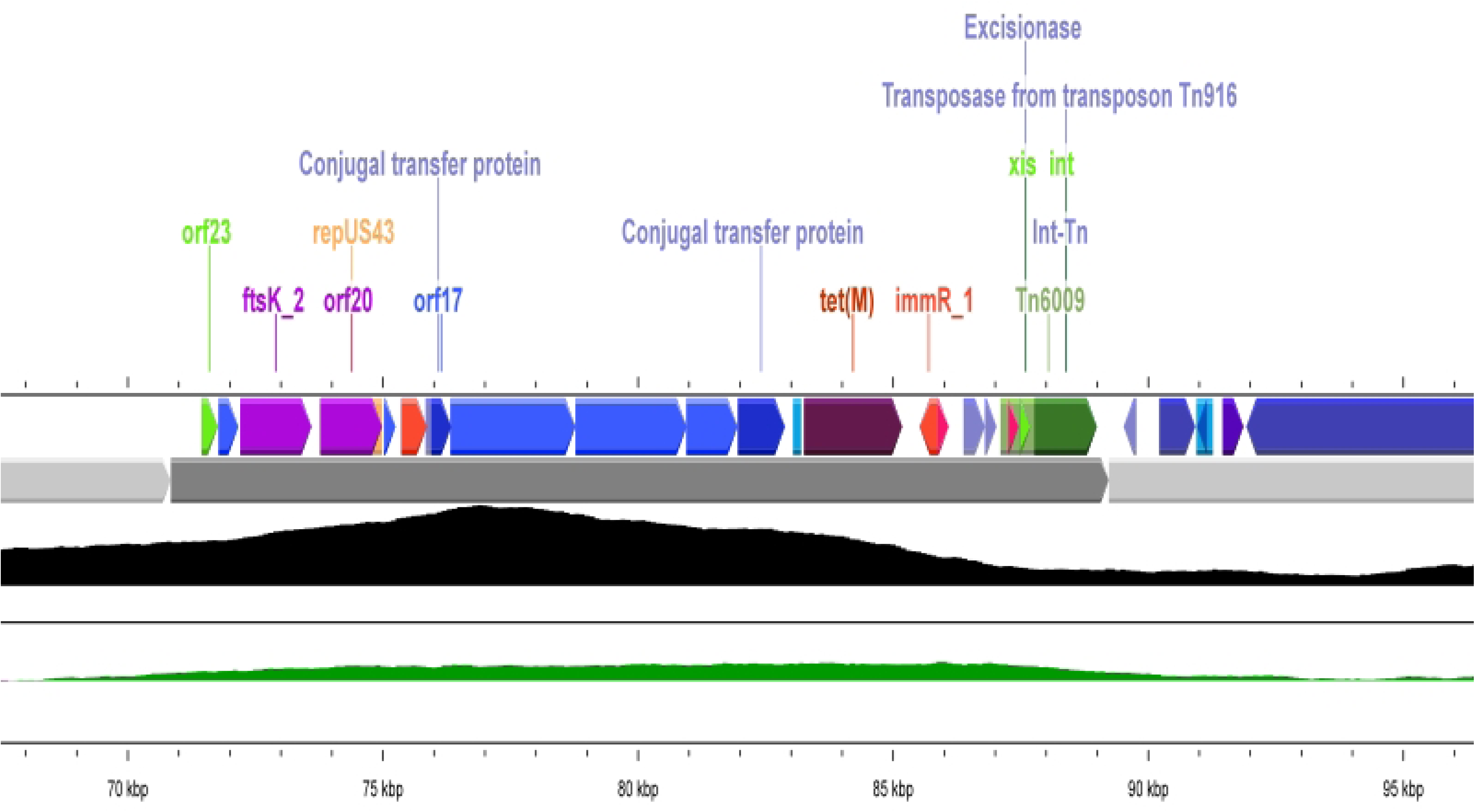
Genomic region of *S. aureus* isolate ERR4692243 showing a partial sequence of the plasmid *repUS43* integrated into the chromosome, located adjacent to open reading frames (ORFs) encoding conjugal transfer proteins and in close proximity to the tetracycline resistance gene *tet(M)*. The region also harbors transposon Tn6009, including its *xis* (excisionase) and *int* (integrase) units, positioned near *repUS43* and *tet(M)*. This physical linkage of plasmid, transposon, and antimicrobial resistance determinants underscores the potential for coordinated horizontal gene transfer and co-selection of resistance traits.

#### Transposons and insertion sequences

Three distinct transposable elements were detected across the genomes. Tn6009 was present in two assemblies, carrying the Int-Tn gene encoding a transposase derived from the Tn916 element (Figure 3). The Tn552 and Tn554 transposons were each found in a single genome. Thirty-one insertion sequences (IS) were identified in 23/40 assemblies (57.5%), with ISSau8 (n = 8) and ISSau3 (n = 6) being the most common. Notably, IS257, a member of the IS26 family that encodes the transposase tnp257, was also detected within an SCCmec element (Fig 4).

**Fig 4:**
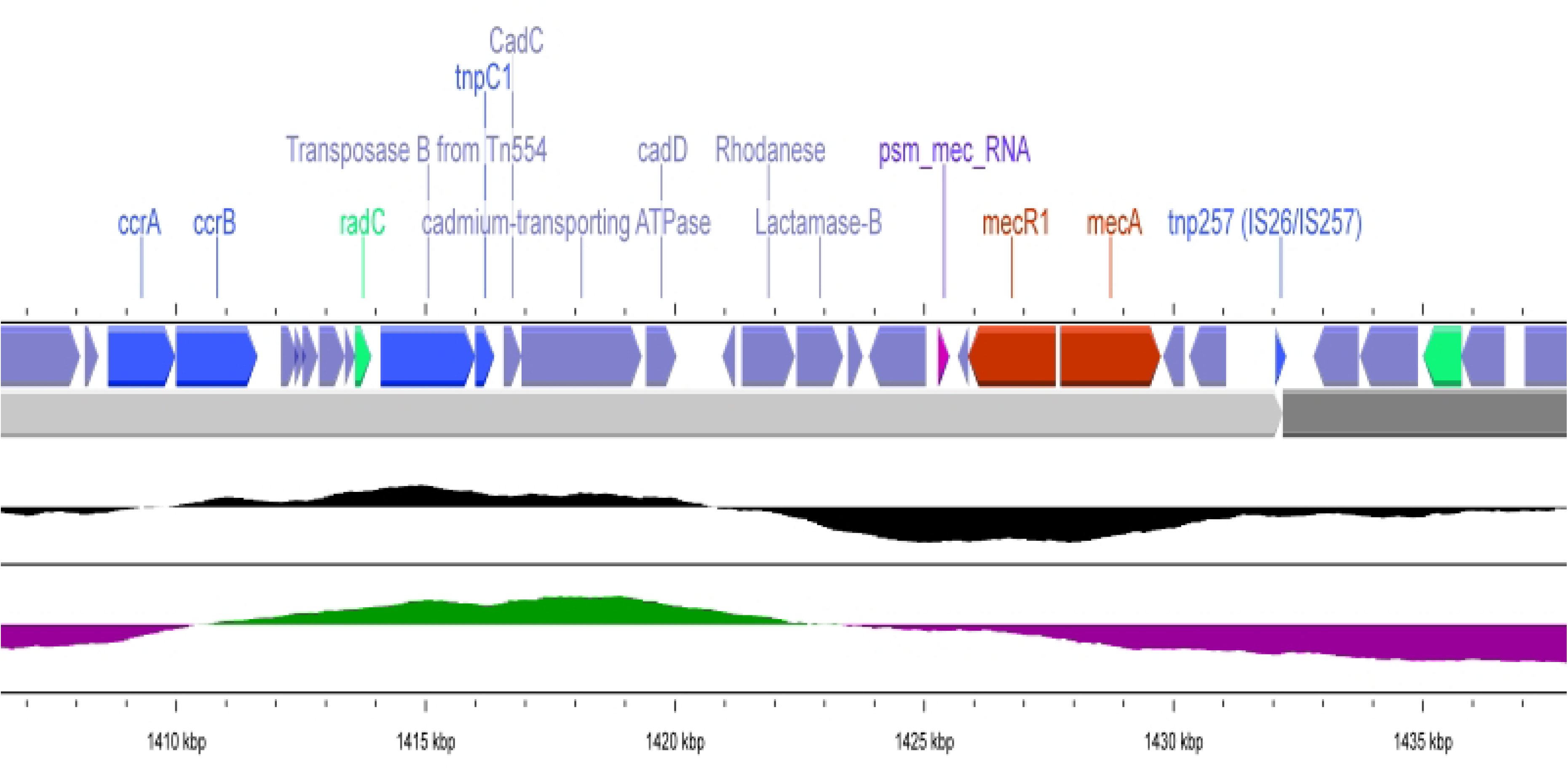
Genomic region of *S. aureus* isolate ERR4692259 depicting the identified SCC*mec* type III (3A) element. The cassette displays the characteristic architecture of SCC*mec* III, flanked upstream by the insertion sequence IS257/tnp257 and downstream by cassette chromosome recombinase genes (*ccrA*/*ccrB*). The core region harbors the methicillin resistance determinant *mecA* together with its regulatory genes *mecI* and *mecR1*. Additional resistance and accessory genes, including *blaZ* (β-lactamase), *cadC* (cadmium resistance transcriptional regulator), and *cadD* (cadmium efflux protein), were also present, highlighting the multifunctional nature of the cassette in conferring resistance to both antibiotics and heavy metals.

#### SCC*mec* cassettes

The *mecA* gene was identified in four genomes, classifying 10% of isolates as methicillin-resistant *S. aureus* (MRSA). Isolates ERR4692243 and ERR4692259 (spa type t37) carried SCC*mec* type IV (2B&5) and SCC*mec* type III (3A), respectively (Fig 1). Other *mecA*-positive isolates ERR4692269 (spa type t1476) and ERR4692272 (spa type t355), lacked a complete SCC*mec* structure but contained SCCmec-associated components (ccr4 and ccr5 complexes, respectively). The SCC*mec* elements we detected not only harbored *mecA* (encoding PBP2a for β-lactam resistance) but also carried *blaZ* (β-lactamase, hydrolyzing penicillins), *cadD,* and *cadC* (cadmium resistance, efflux, and regulatory proteins), along with hallmark mobile components such as insertion sequence IS257 and transposon Tn554 (Fig 4).

#### Prophages

We identified 147 prophage sequences in the 40 *S. aureus* genomes. These were grouped into three categories by PHASTER according to their completeness (47 intact, 40 questionable, and 60 incomplete). Only intact prophages were considered for manual annotation and are indicated in Figure 1. The lengths of the intact prophages ranged from 16.0 to 64.0 Kb with a GC content between 33.3% and 36.95%. The most frequently identified phages were *Staphylococcus* PHAGE_phiJB (n = 13), *Staphylococcus* PHAGE_96 (n = 9), and *Staphylococcus* PHAGE_phi2958PVL (n = 9), all belonging to the Siphoviridae family. The 47 intact phages were manually annotated to detect antimicrobial resistance genes (ARG) and virulence factors (Table 1 and Fig 5).

**Fig 5:**
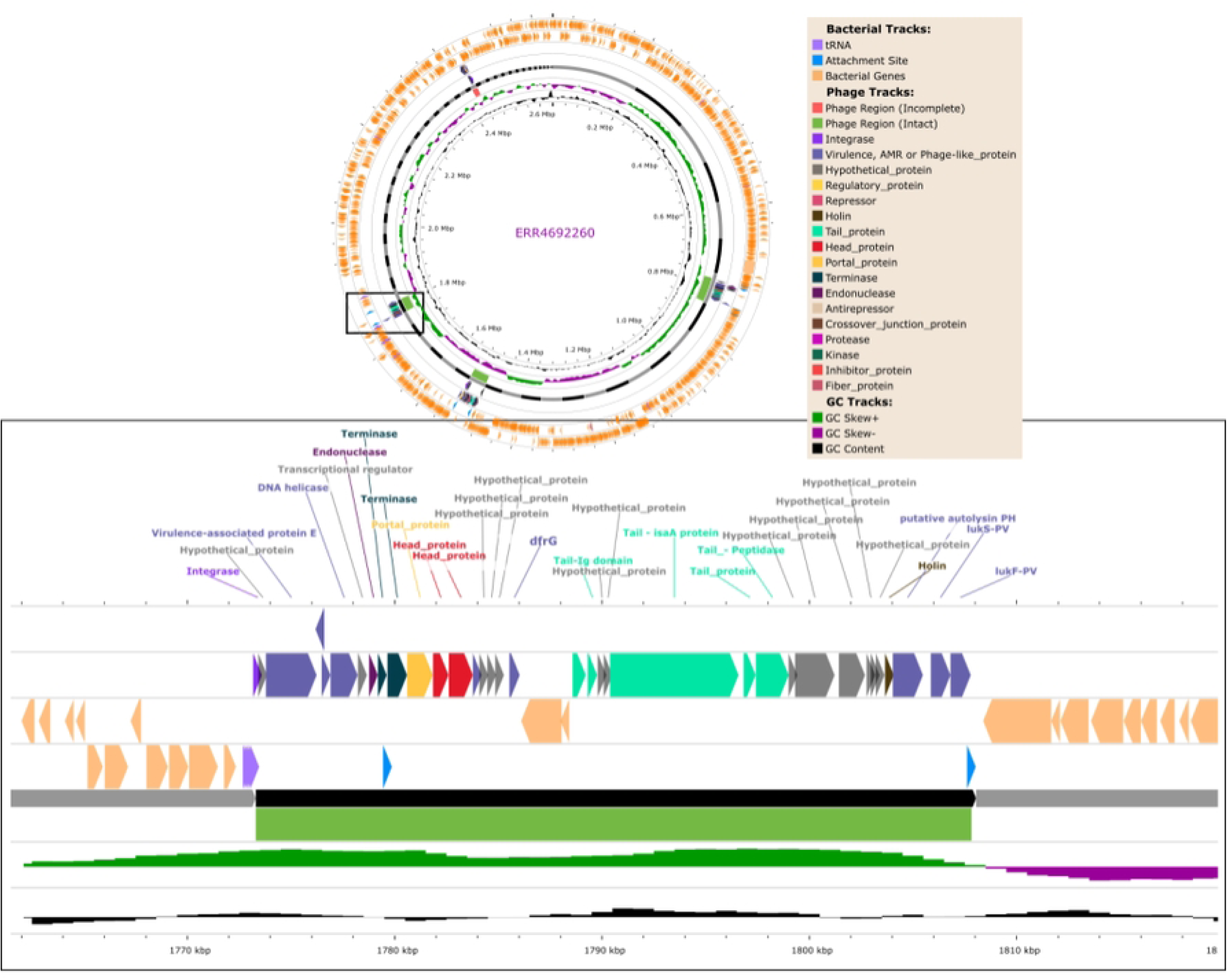
Prophage landscape of *S. aureus* isolate ERR4692260, revealing four distinct prophage regions identified by PHASTER—three intact and one incomplete. One of the intact regions, classified as *PHAGE_Staphy_phi2958PVL*, displays hallmark features of a functional prophage, including genes encoding integrase, capsid/head, portal, tail, and holin proteins. Manual curation further identified key accessory genes, such as the antimicrobial resistance gene *dfrG* and the leukocidin toxin genes *lukF-PV* and *lukS-PV*, highlighting the potential of this prophage to contribute to both resistance and virulence in the host strain.

**Table 1:**
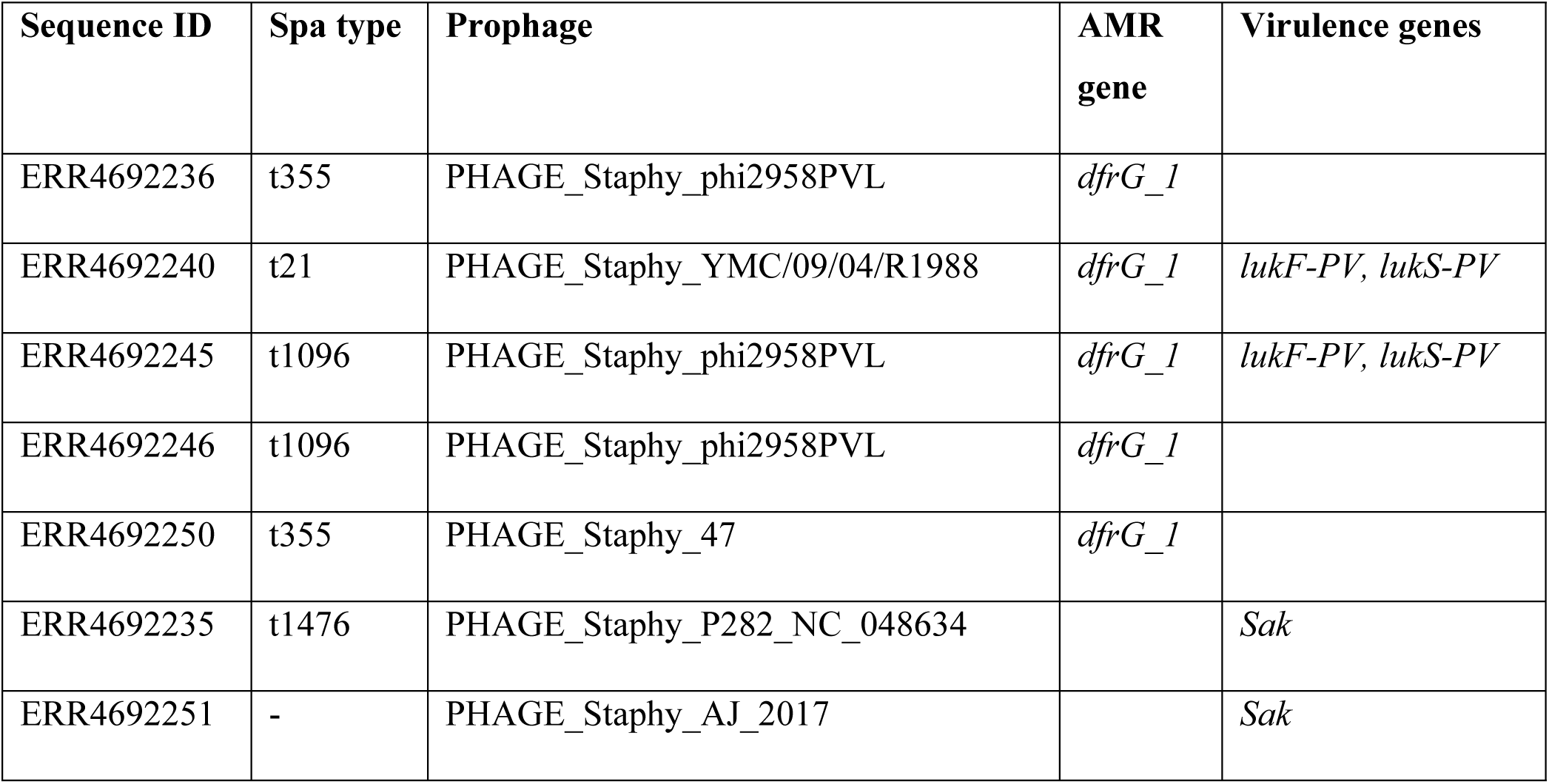
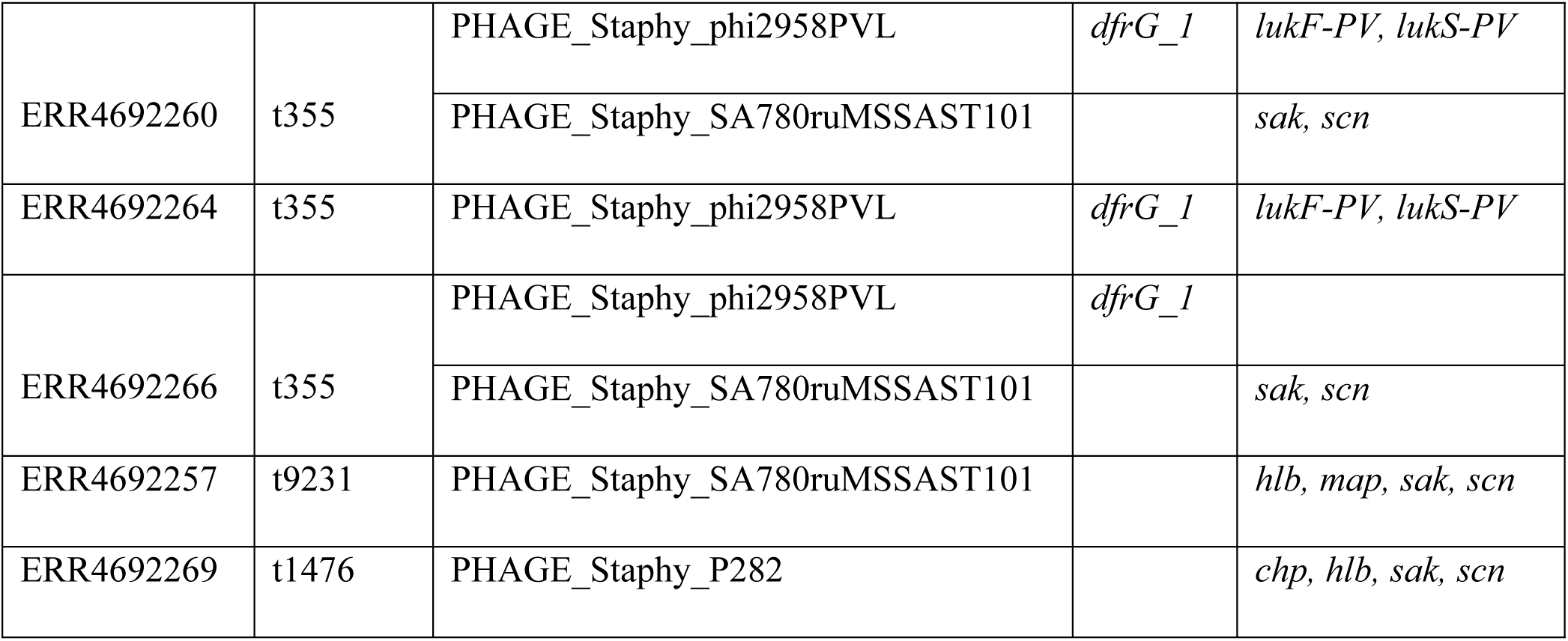
AMR and virulence-encoding genes in prophages by manual annotation.

#### Genomic Islands

Genomic island (GI) analysis across all 40 *Staphylococcus aureus* genome assemblies, using IslandViewer4, revealed a total of 191 genomic islands. Each genome harbored at least one island, with the highest counts observed in isolates ERR4692243 (n = 15), ERR4692259 (n = 13), and ERR4692237 (n = 11). These genomic islands encoded a diverse array of functional genes, notably those associated with virulence, immune evasion, and antimicrobial resistance (Fig 6). Core virulence factors included *nuc* (thermonuclease), *sak* (staphylokinase), *tst* (toxic shock syndrome toxin), and *lukF-PV/lukS-PV* (Panton-Valentine leukocidin), the latter of which is strongly implicated in severe necrotizing infections. Several immune evasion-associated genes were also identified within GIs, including *vapE* (virulence-associated protein E), *hmp* (flavohemoprotein for nitric oxide detoxification), *hemolysin III* (membrane-damaging cytotoxin), *EF-Tu* and *tuf* (elongation factor Tu, moonlighting in adhesion and immune evasion), *enolase* (surface-bound plasminogen-binding protein), *cold shock proteins*, and *AhpC* (alkyl hydroperoxide reductase, involved in oxidative stress defense). In addition, antimicrobial and metal resistance genes were frequently embedded in these regions. These included *blaZ* (β-lactamase), *ermC* (macrolide resistance), *tet* (tetracycline resistance), *sepA* (multidrug efflux), *dfrG* and *dfrC* (trimethoprim resistance), *lmrs* (linezolid resistance), as well as c*adD* (cadmium resistance), *copG* (copper efflux regulation), and *mer* (mercuric resistance). (Fig 6)

**Fig 6:**
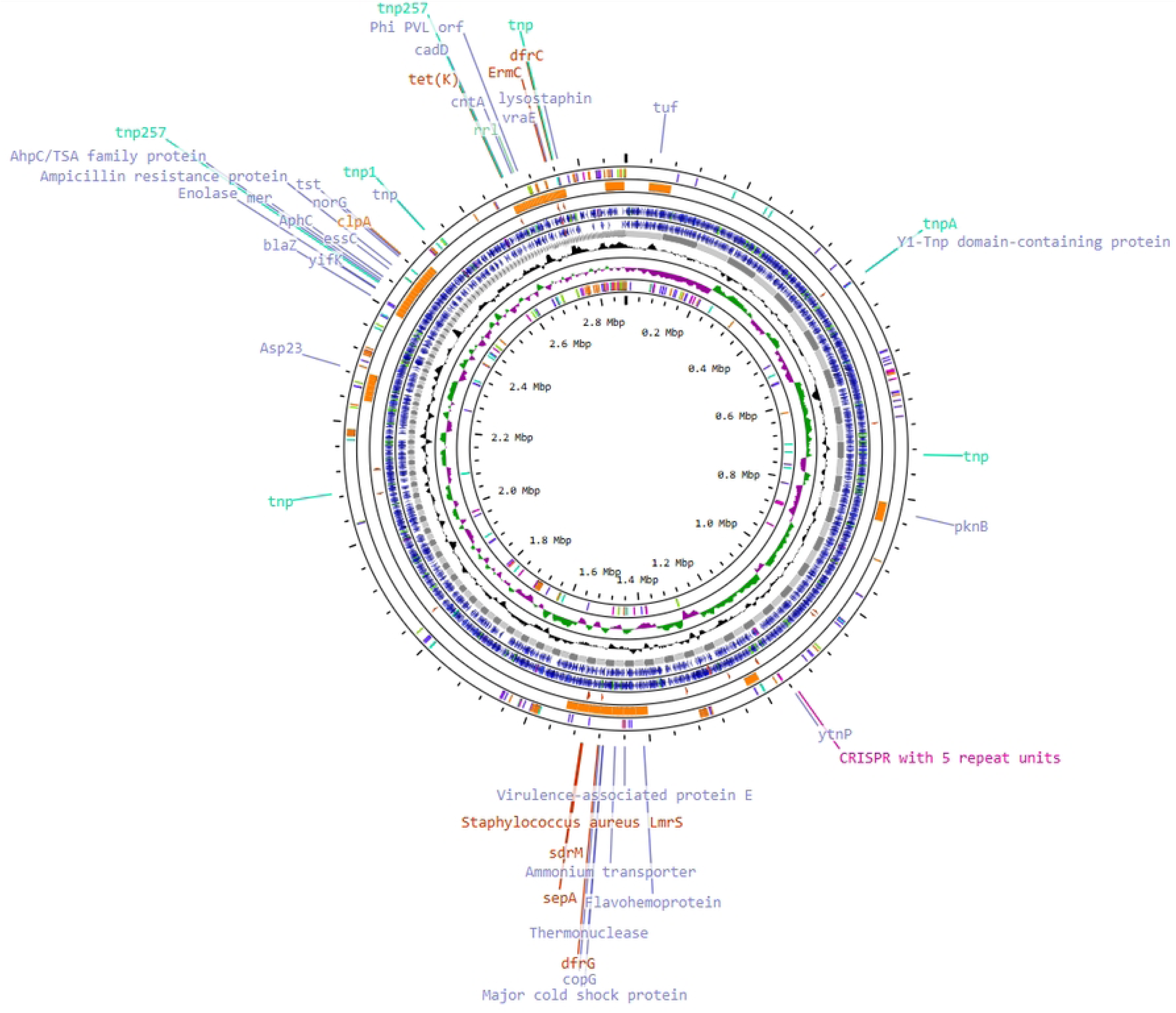
Circularized genome of *S. aureus* isolate ERR4692263, illustrating genomic islands and associated antimicrobial resistance (AMR) and virulence genes. Genomic islands were identified using IslandViewer4, which integrates sequence composition-based (SIGI-HMM, IslandPath-DIMOB) and comparative genomics approaches (IslandPick). From the outermost to the innermost ring:Ring 1 displays the mobile genetic elements as predicted by mobile genetic finder (+ strand); Ring 2 displays predicted genomic islands; Ring 3 shows AMR genes annotated via CARD (Comprehensive Antibiotic Resistance Database); Ring 4 presents functional gene annotations from Bakta; Ring 5 includes key genomic features such as virulence factors; Ring 6 shows the genomic backbone across assembled contigs; Ring 7 plots GC content variation; Ring 8 illustrates GC skew; and Ring 9 shows mobile genetic elements (-strand). The figure highlights the genomic architecture and localized clustering of resistance and virulence determinants within predicted genomic islands.

## Discussion

This study is the first to characterize *S. aureus* spa types circulating in FPRRH, western Uganda. We identified 11 spa types, with t355 being the most prevalent (35%, 14/40). Previously, t355 was primarily associated with obstetrics and gynecology wards at Mulago Hospital in central Uganda [17]. Our findings extend this spa type’s distribution to western Uganda, reinforcing its prominence across the country and also supporting previous reports from eastern Uganda [2] and several counties in Kenya [18]. Notably, our study provided higher resolution typing compared to earlier MLST-based work on the same isolates [19], identifying 11 spa types versus 9 sequence types, underscoring the greater discriminatory power of spa-typing for *S. aureus* epidemiology [20].

We identified two SCC*mec* types: type III (HA-MRSA) and type IV (CA-MRSA), confirming the coexistence of these elements previously reported in eastern Uganda [21]. MRSA genomes belonged to spa types t37, t355, and t1476, with t37 consistently observed in multiple Ugandan studies [2, 21], suggesting its potential endemicity. The *mecA* gene, encoding PBP2a for β-lactam resistance, was detected in four genomes (10%); however, complete SCC*mec* structures were only present in ERR4692243 (spa type t37, SCC*mec* IV [2B&5]) and ERR4692259 (SCC*mec* III [3A]) (Figures 1 and 2). The remaining two *mecA*-positive isolates (ERR4692269, spa type t1476; ERR4692272, spa type t355) lacked full cassettes but contained key structural components (*ccr4* and *ccr5* complexes, respectively), consistent with previous reports of partial or degraded SCCmec elements [50]. Such genomic streamlining may reflect adaptive strategies to mitigate the fitness costs associated with maintaining large SCC*mec* structures while retaining essential resistance determinants.

Beyond *mecA*, SCC*mec* regions in this study carried additional resistance and accessory genes, notably *blaZ* (β-lactamase; hydrolyzes penicillins), cadD, and *cadC* (cadmium efflux pump and transcriptional regulator), alongside hallmark mobile elements such as IS257 and transposons. IS257, a member of the IS26 family, is known to mediate SCC*mec* structural rearrangements, mobilization of *mecA*, and plasmid–chromosome recombination events and has been implicated in the co-mobilization of metal and antimicrobial resistance genes [59, 60, 61]. The co-localization of antimicrobial and metal resistance determinants within SCC*mec* underscores their multifunctional role in adaptation to both antibiotic and heavy metal stressors, while the association with integrases and insertion sequences affirms their potential for mobility and structural evolution within the *S. aureus* mobilome.

None of the MRSA genome sequences carrying SCC*mec* elements harbored *lukPV* genes encoding Panton-Valentine leukocidin (PVL). In contrast, four MSSA genomes in this study did carry PVL genes, which were located on two distinct intact prophages: phi2958PVL and YMC/09/04/R1988. Among these, phi2958PVL accounted for 75% of all detected PVL genes. Notably, these prophages also harbored the trimethoprim resistance gene *dfrG*, highlighting a concerning co-localization of virulence and resistance determinants within the same mobile genetic elements. This finding aligns with prior studies that have documented Siphoviridae phages carrying both *lukPV* and resistance genes in *S. aureus*, contributing to the emergence of successful clones such as CC80 and CC30, which are widely associated with PVL-positive CA-MRSA and have been implicated in epidemics across Europe and Africa [51, 52]. Specifically, *dfrG* has been reported on phage-plasmid hybrids and chromosomal islands and frequently co-occurs with insertion sequences, which facilitate integration and mobilization [53]. Our detection of *dfrG* on phages is consistent with recent observations of ARG-carrying prophages in *S. sciuri* and *S. aureus*, including those bearing aminoglycoside resistance (*aacA-aphD*) and *dfrA* [54], further supporting the role of phages in capturing and disseminating ARGs across species.

Our observation that several PVL-encoding prophages also carried *dfrG* highlights the emerging role of phages as composite mobile elements linking virulence and resistance in *S. aureus*. This co-localization mirrors findings from other studies, where PVL-positive prophages have been shown to harbor antibiotic resistance genes, including *tetK, ermC*, and *blaZ*, supporting the view that temperate phages can act as hybrid vectors of both pathogenicity and antimicrobial resistance [55, 56]. Such dual-function phages not only enhance the fitness of *S. aureus* strains under selective pressure such as antibiotics but may also accelerate the spread of high-risk clones in community and hospital settings.

These findings underscore the importance of phage-mediated horizontal gene transfer in shaping the resistome and virulome of *S. aureus*. The integration of *dfrG* and *lukPV* within the same mobile element may provide a selective advantage, promoting the emergence of virulent, drug-resistant MSSA lineages, which may be overlooked in MRSA-focused surveillance. This has important implications for regional surveillance and highlights the need for genome-based diagnostics that can capture such complex genomic architectures.

Mobile antibiotic resistance genes (ARGs) are frequently associated with plasmids, which are major vectors of horizontal gene transfer in *S. aureus*. Although plasmid DNA was not physically extracted in the original study [19], our whole-genome sequence analysis revealed 74 plasmid-derived sequences, with *repUS43*, *rep7a,* and *rep16* among the most common replicons. Notably, *repUS43* carried a MOBT-type relaxase, a signature of mobilizable plasmids capable of mediating transfer of integrative and conjugative elements across Firmicutes [22]. While such plasmids are often described as cryptic due to the absence of obvious phenotypic traits, increasing evidence shows they serve as moldable vectors, poised to acquire and disseminate adaptive traits, including antimicrobial resistance [23]. Interestingly, all plasmid-derived sequences in our dataset were integrated within chromosomal contigs rather than existing as extrachromosomal elements. Chromosomal capture of plasmid material has been described in clinical S. aureus populations and is increasingly recognized as an important mechanism that both stabilizes accessory genes and creates reservoirs for later mobilization [57, 58]. This chromosomal capture likely results from recombination or transposition events facilitated by mobile elements such as recombinases or transposases, allowing plasmid cargo to be stably inherited vertically while retaining potential for future mobilization. The proximity of these integrated plasmid sequences to ORFs encoding conjugal transfer proteins, AMR determinants such as *tet(K)*, and other MGEs supports the notion of historical or ongoing horizontal gene exchange.

Three distinct transposons were identified, including Tn6009, a derivative of Tn916, which was found in two genomes. This element is particularly notable for carrying the *Int-Tn* gene encoding a Tn916-family transposase and for its established linkage to the *S. aureus mer* operon (conferring mercury resistance) and *tet(M)* tetracycline resistance [25]. Historically, Tn6009 was first described in Gram-positive bacteria in Portugal in 1997 and later detected in *Klebsiella pneumoniae* from Nigeria (2002–2003) [24], with more recent detection in *Enterococcus faecalis* in South Africa in 2021 [26]. In our study, Tn6009 was identified in Uganda for the first time, associated with *tet(M)* as previously reported in other settings. The conservation of this *tet(M)* association across diverse hosts and regions suggests a stable genetic linkage, potentially maintained by co-selection pressures from tetracycline use. Given its broad host range, capacity for ARG acquisition, and confirmed presence in geographically distant regions, Tn6009 may represent an underappreciated driver of tetracycline resistance dissemination in African *S. aureus* populations.

We also detected 31 insertion sequences in 23 genomes (57.5%), with ISSau8 and ISSau3 being the most frequent. Of particular note was IS257, a member of the IS26 family, identified within the SCC*mec* type III element in our dataset. IS257 has been well documented as a key driver of SCC*mec* structural rearrangements, mobilization of the *mecA* gene, and recombination events between plasmids and the chromosome in *S. aureus* [60]. It has also been implicated in the co-mobilization of additional resistance determinants, including β-lactamases (*blaZ*), cadmium-resistance genes (*cadD, cadC*), and aminoglycoside-modifying enzymes [60, 61]. The detection of IS257 within SCC*mec* type III here underscores the dynamic and composite nature of these elements, which not only carry *mecA* but also serve as hotspots for the accumulation of diverse adaptive genes through IS-mediated recombination.

Closer examination of the *S. aureus* genomes revealed a high density of virulence and resistance genes embedded within genomic islands (GIs), underscoring their critical role in bacterial adaptation and pathogenicity. Notably, these islands harbored genes linked to virulence, immune evasion, and antimicrobial resistance. Core virulence determinants included *nuc* (thermonuclease), *sak* (staphylokinase), *tst* (toxic shock syndrome toxin), and *lukF-PV*/*lukS-PV* (Panton-Valentine leukocidin), the latter of which is strongly implicated in necrotizing pneumonia and severe skin infections. Several GIs also encoded factors associated with immune evasion and stress resistance, such as *vapE* (virulence-associated protein E), *hmp* (flavohemoprotein involved in nitric oxide detoxification), hemolysin III (a membrane-damaging cytotoxin), *EF-Tu* and *tuf* (elongation factor Tu, which moonlights in adhesion and evasion of host defenses), enolase (a surface-bound plasminogen-binding protein), cold shock proteins, and *AhpC* (alkyl hydroperoxide reductase involved in oxidative stress response). The GIs also harbored genes conferring resistance to antibiotics and heavy metals. These included *blaZ* (β-lactamase), *ermC* (macrolide resistance), *tet* (tetracycline resistance), *sepA* (multidrug efflux pump), *dfrG* and *dfrC* (trimethoprim resistance), and *lmrs* (linezolid resistance). Additionally, metal resistance genes such as *cadD* (cadmium resistance), *copG* (copper efflux regulation), and *mer* (mercuric resistance) were commonly detected, suggesting adaptation to both clinical and environmental pressures.

Importantly, the mobility of these GIs was supported by the presence of integrases, transposases, insertion sequences (notably IS257), and occasional CRISPR-associated genes, as well as GC content skew relative to the host genome, which are classical signatures of horizontal acquisition. Such features indicate that these GIs are not static genetic cargo but dynamic, mobilizable regions capable of inter-strain transfer. Comparable patterns have been documented in *Helicobacter pylori* from Kenya [27] and *Vibrio cholerae* from Tanzania [28].

## Conclusions

*S. aureus* isolates from FPRRH in western Uganda carry a rich array of mobile genetic elements (MGEs), including genomic islands, prophages, plasmid-derived sequences, transposons, and SCC*mec* cassettes. These MGEs harbored key virulence factors (*lukPV, tst, nuc*) and resistance genes (*mecA, dfrG, tet*), often co-localized within the same mobile units, particularly phages, highlighting their role in both pathogenesis and drug resistance. The predominance of spa type t355 and detection of both hospital- and community-associated MRSA lineages suggest ongoing local evolution and potential transmission overlap. The integration of MGEs into chromosomal regions, alongside signatures of mobility such as integrases and insertion sequences, reveals an active and adaptable mobilome. These findings underscore the need for targeted genomic surveillance to track and mitigate the spread of high-risk *S. aureus* clones in Uganda.

## Methods

### Study design and setting

This was a cross-sectional study conducted at the Genomics and Molecular Biology Laboratory of Makerere University College of Health Sciences, and the African Centre of Excellence in Bioinformatics & Data Intensive Sciences, Infectious Diseases Institute, Makerere University College of Health Sciences. The study used a total of 40 *S. aureus* whole genome sequences previously characterized by Ackers-Johnson *et al.* [19]; these sequences represent the same number of isolates, i.e., 40 clinically relevant *S. aureus* isolates from skin wounds, urinary tract, or bloodstream infections from patients at FPRRH, Western Uganda, that were collected and phenotypically characterized by the FPRRH microbiology laboratory between 2017 and 2019.

### Retrieval of the whole genome sequences

A total of 40 whole genome sequences of the *S. aureus* isolates described above were downloaded from the European Nucleotide Archive (https://www.ebi.ac.uk/ena) by searching and navigating to the project accession number PRJEB40863.

### Draft genome assembly and annotation

Illumina sequencing adapters and bad reads were trimmed from the raw downloaded sequences using fastp [47] and assembled using SPADES v3.9.1 [31]. The trimmed reads were mapped to the *S. aureus* reference genome NCTC 8325 using Burrows-Wheeler Aligner software (BWA-MEM) [32] for reference-based genome assembly. The resulting draft assemblies were then annotated using PROKKA v 1.13.3 [33].

### In-silico identification of spa types

In silico spa genotyping was achieved using spaTyper version 1.0 with default settings [34]. This tool used spa sequences database based on the repeat sequences and repeat orders from the spa typing website http://www.spaserver.ridom.de/

### In silico identification of mobile genetic elements

Plasmids were predicted from the assembled genomes using PlasmidFinder v2.1 [35,36] with the Gram-positive database, applying thresholds of ≥95% sequence identity and ≥60% coverage. PHASTEST (PHAge Search Tool with Enhanced Sequence Translation) was employed to detect and annotate prophage sequences [37, 38, 42]. This web-based platform enables rapid identification, annotation, and visualization of prophages within bacterial chromosomes and plasmids, with intact prophages defined by a quality score >90.

Insertion sequences and transposons were identified using MobileElementFinder v1.0.3 under default parameters [39]. SCCmecFinder v1.2 [36, 40, 41] was used to detect SCCmec elements, applying thresholds of ≥90% identity and ≥60% minimum length against the reference database. Genomic islands were predicted using IslandViewer4 [43], which integrates three complementary methods: IslandPick, SIGI-HMM [44], and IslandPath-DIMOB [45] to detect genomic and pathogenicity islands.

Further screening, annotation, and visualization of mobile genetic elements (MGEs) were conducted using ProkSee [46], an expert genome assembly, annotation, and visualization system that supports interactive circular and linear genome mapping. Within ProkSee, Bakta [48] was applied for rapid and standardized annotation of genomes and plasmids; MobileOG-db [49] was used to identify protein families mediating integration/excision, replication/recombination/repair, stability/defense, and transfer of bacterial MGEs and phages, along with associated transcriptional regulators; and the Comprehensive Antibiotic Resistance Database (CARD) [62] was employed for annotation of antimicrobial resistance genes, their products, and associated phenotypes.

## Ethical considerations

Ethical approval was received from the Makerere University School of Biomedical Sciences Research & Ethics Committee (#SBS-2022-266); a waiver of informed consent from the participants from whom organisms that were sequenced were isolated was granted by the Makerere SBS Ethics Committee.

## Data availability

The genome sequences included in this study are available from ENA https://www.ebi.ac.uk/ena under project number PRJEB40863.

## Authors contributions

Conceptualization: MLN, EK, KE, KP, FY, BW, MR, MWJ, KL, HS, FAK, GAJ, CJ, DPK Bioinformatics Analysis: MLN, EK, KE, KP, FY, BW, MWJ, KL, HS, FAK, GAJ, CJ, DPK Manuscript writing: MLN, DPK, EK, KP, HS All authors read and approved the final version of the manuscript.

## Acknowledgements

We gratefully acknowledge the management and staff of the Genomics, Molecular, and Immunology Laboratories in the Department of Immunology and Molecular Biology, Makerere University College of Health Sciences, for the technical and moral support that made this work a success.

Research reported in this publication was supported by the Fogarty International Center of the National Institutes of Health under Award Numbers U2RTW010672 (Nurturing Genomics and Bioinformatics Research Capacity in Africa program – BRECA) and U2RTW012116 (Makerere University Data Science Research Training to Strengthen Evidence-Based Health Innovation, Intervention and Policy – MakDARTA), and by the Cambridge-Africa Program of the University of Cambridge. The content is solely the responsibility of the authors and does not necessarily represent the official views of the funders.

## References

1. Kateete, D. P. et al. High prevalence of methicillin-resistant Staphylococcus aureus in the surgical units of Mulago Hospital in Kampala, Uganda. BMC Res Notes 4, 326 (2011).

2. Kateete, D. P., et al. Nasopharyngeal carriage, spa types, and antibiotic susceptibility profiles of Staphylococcus aureus from healthy children less than 5 years old in Eastern Uganda. BMC Infect Dis 19, 1023 (2019).

3. Gajdács, M. The Continuing Threat of Methicillin-Resistant Staphylococcus aureus. Antibiotics 8, 52 (2019).

4. World Health Organization. Antimicrobial resistance. (2020).

5. Kobayashi, S. D., Musser, J. M. & DeLeo, F. R. Genomic Analysis of the Emergence of Vancomycin-Resistant Staphylococcus aureus. mBio 3, e00170–12 (2012).

6. Otto, M. Staphylococcus aureus toxins. Current Opinion in Microbiology 17, 32–37 (2014).

7. McCarthy, A. J., et al. Extensive Horizontal Gene Transfer during Staphylococcus aureus Co-colonization In Vivo. Genome Biology and Evolution 6, 2697–2708 (2014).

8. Holden, M. T. G., et al. Complete genomes of two clinical *Staphylococcus aureus* strains: Evidence for the rapid evolution of virulence and drug resistance. Proc. Natl. Acad. Sci. U.S.A. 101, 9786–9791 (2004).

9. Malachowa, N. & DeLeo, F. R. Mobile genetic elements of Staphylococcus aureus. Cell. Mol. Life Sci. 67, 3057–3071 (2010).

10. Ferreira, C. et al. Clonal Lineages, Antimicrobial Resistance, and PVL Carriage of Staphylococcus aureus Associated to Skin and Soft-Tissue Infections from Ambulatory Patients in Portugal. Antibiotics 10, 345 (2021).

11. Bukowski, M. et al. Prevalence of Antibiotic and Heavy Metal Resistance Determinants and Virulence-Related Genetic Elements in Plasmids of Staphylococcus aureus. Front. Microbiol. 10, 805 (2019).

12. Jani, M., Sengupta, S., Hu, K. & Azad, R. K. Deciphering pathogenicity and antibiotic resistance islands in methicillin-resistant *Staphylococcus aureus* genomes. Open Biol. 7, 170094 (2017).

13. Nemerovski, C. W. & Klein, K. C. Community-Associated Methicillin-Resistant Staphylococcus aureus in the Pediatric Population. The Journal of Pediatric Pharmacology and Therapeutics 13, 212–225 (2008).

14. McCarthy, A. J., Witney, A. A. & Lindsay, Jodi. A. Staphylococcus aureus Temperate Bacteriophage: Carriage and Horizontal Gene Transfer is Lineage Associated. Front. Cell. Inf. Microbio. 2, (2012).

15. Jamrozy, D. et al. Evolution of mobile genetic element composition in an epidemic methicillin-resistant Staphylococcus aureus: temporal changes correlated with frequent loss and gain events. BMC Genomics 18, 684 (2017).

16. Inzaule, S. C., Tessema, S. K., Kebede, Y., Ogwell Ouma, A. E. & Nkengasong, J. N. Genomic-informed pathogen surveillance in Africa: opportunities and challenges. The Lancet Infectious Diseases 21, e281–e289 (2021).

17. Seni, J., et al. Molecular Characterization of Staphylococcus aureus from Patients with Surgical Site Infections at Mulago Hospital in Kampala, Uganda. PLoS ONE 8, e66153 (2013).

18. Nyasinga, J. et al. Epidemiology of Staphylococcus aureus< Infections in Kenya: Current State, Gaps, and Opportunities. OJMM 10, 204–221 (2020).

19. Ackers-Johnson, G., et al. Antibiotic resistance profiles and population structure of disease-associated Staphylococcus aureus infecting patients in Fort Portal Regional Referral Hospital, Western Uganda. Microbiology 167, (2021).

20. Cheşcă, A. Evaluation of sequence-based typing methods (spa and MSLT) for clonal characterization of Staphylococcus aureus. Acta Medica Mediterranea (2016) doi:10.19193/0393-6384_2016_6_173.

21. Kateete, D. P. et al. CA-MRSA and HA-MRSA coexist in community and hospital settings in Uganda. Antimicrob Resist Infect Control 8, 94 (2019).

22. Ramachandran, G., et al. Discovery of a new family of relaxases in Firmicutes bacteria. PLoS Genet 13, e1006586 (2017).

23. Attéré, S. A., Vincent, A. T., Paccaud, M., Frenette, M. & Charette, S. J. The Role for the Small Cryptic Plasmids As Moldable Vectors for Genetic Innovation in Aeromonas salmonicida subsp. salmonicida. Front. Genet. 8, 211 (2017).

24. Soge, O. O., Beck, N. K., White, T. M., No, D. B. & Roberts, M. C. A novel transposon, Tn6009, composed of a Tn916 element linked with a Staphylococcus aureus mer operon. Journal of Antimicrobial Chemotherapy 62, 674–680 (2008).

25. Roberts, A. P. & Mullany, P. Tn *916*-like genetic elements: a diverse group of modular mobile elements conferring antibiotic resistance. FEMS Microbiol Rev 35, 856–871 (2011).

26. Mbanga, J. et al. Genomic Analysis of Enterococcus spp. Isolated From a Wastewater Treatment Plant and Its Associated Waters in Umgungundlovu District, South Africa. Front. Microbiol. 12, 648454 (2021).

27. Mwangi, C., et al. Whole Genome Sequencing Reveals Virulence Potentials of Helicobacter pylori Strain KE21 Isolated from a Kenyan Patient with Gastric Signet Ring Cell Carcinoma. Toxins 12, 556 (2020).

28. Hounmanou, Y. M. G. et al. Genomic insights into Vibrio cholerae O1 responsible for cholera epidemics in Tanzania between 1993 and 2017. PLoS Negl Trop Dis 13, e0007934 (2019).

29. Liu, B. et al. In silico Evolution and Comparative Genomic Analysis of IncX3 Plasmids Isolated From China Over Ten Years. Front. Microbiol. 12, 725391 (2021).

30. Bertelli, C. & Brinkman, F. S. L. Improved genomic island predictions with IslandPath-DIMOB. Bioinformatics 34, 2161–2167 (2018).

31. Bankevich, A. et al. SPAdes: A New Genome Assembly Algorithm and Its Applications to Single-Cell Sequencing. Journal of Computational Biology 19, 455–477 (2012).

32. Li, H. & Durbin, R. Fast and accurate short read alignment with Burrows-Wheeler transform. Bioinformatics 25, 1754–1760 (2009).

33. Seemann, T. Prokka: rapid prokaryotic genome annotation. Bioinformatics 30, 2068–2069 (2014).

34. Bartels, M. D., et al. Comparing Whole-Genome Sequencing with Sanger Sequencing for *spa* Typing of Methicillin-Resistant Staphylococcus aureus. J Clin Microbiol 52, 4305–4308 (2014).

35. Carattoli, A., et al. In Silico Detection and Typing of Plasmids using PlasmidFinder and Plasmid Multilocus Sequence Typing. Antimicrob Agents Chemother 58, 3895–3903 (2014).

36. Camacho, C., et al. BLAST+: architecture and applications. BMC Bioinformatics 10, 421 (2009).

37. Zhou, Y., Liang, Y., Lynch, K. H., Dennis, J. J. & Wishart, D. S. PHAST: A Fast Phage Search Tool. Nucleic Acids Research 39, W347–W352 (2011).

38. Arndt, D., et al. PHASTER: a better, faster version of the PHAST phage search tool. Nucleic Acids Res 44, W16–W21 (2016).

39. Johansson, M. H. K. et al. Detection of mobile genetic elements associated with antibiotic resistance in *Salmonella enterica* using a newly developed web tool: MobileElementFinder. Journal of Antimicrobial Chemotherapy 76, 101–109 (2021).

40. Kondo, Y., et al. Combination of Multiplex PCRs for Staphylococcal Cassette Chromosome *mec* Type Assignment: Rapid Identification System for *mec*, *ccr*, and Major Differences in Junkyard Regions. Antimicrob Agents Chemother 51, 264–274 (2007).

41. Classification of Staphylococcal Cassette Chromosome *mec* (SCCmec)): Guidelines for Reporting Novel SCCmecElements. Antimicrobial Agents Chemother 53, 4961–4967 (2009).

42. David S. Wishart, Scott Han, Sukanta Saha, Eponine Oler, Harrison Peters Jason Grant, Paul Stothard, Vasuk Gautam (2023), PHASTEST: Faster than PHASTER, Better than PHAST, Nucleic Acids Research (Web Server Issue), 10.1093/nar/gkad382

43. Bertelli, C. et al. IslandViewer 4: expanded prediction of genomic islands for larger-scale datasets. Nucleic Acids Research 45, W30–W35 (2017).

44. Waack, S. et al. Score-based prediction of genomic islands in prokaryotic genomes using hidden Markov models. BMC Bioinformatics 7, 142 (2006).

45. Hsiao, W., Wan, I., Jones, S. J. & Brinkman, F. S. L. IslandPath: aiding detection of genomic islands in prokaryotes. Bioinformatics 19, 418–420 (2003).

46. Grant JR, Enns E, Marinier E, Mandal A, Herman EK, Chen C, Graham M, Van Domselaar G, and Stothard P, Proksee: in-depth characterization and visualization of bacterial genomes Nucleic Acids Research, 2023, gkad326, 10.1093/nar/gkad326

47. Shifu Chen, Yanqing Zhou, Yaru Chen, Jia Gu; fastp: an ultra-fast all-in-one FASTQ preprocessor, Bioinformatics, Volume 34, Issue 17, 1 September 2018, Pages i884–i890, 10.1093/bioinformatics/bty560.

48. Schwengers O., Jelonek L., Dieckmann M. A., Beyvers S., Blom J., Goesmann A. (2021). Bakta: rapid and standardized annotation of bacterial genomes via alignment-free sequence identification. Microbial Genomics, 7(11). 10.1099/mgen.0.000685

49. Brown CL, Mullet J, Hindi F, Stoll JE, Gupta S, Choi M, Keenum I, Vikesland P, Pruden A, Zhang L. mobileOG-db: a Manually Curated Database of Protein Families Mediating the Life Cycle of Bacterial Mobile Genetic Elements. Appl Environ Microbiol. 2022 Aug 29:e0099122. doi: 10.1128/aem.00991-22. Epub ahead of print. PMID: 36036594.

50. Noto MJ, Fox PM, Archer GL. Spontaneous deletion of the methicillin resistance determinant, mecA, partially compensates for the fitness cost associated with high-level vancomycin resistance in Staphylococcus aureus. Antimicrob Agents Chemother. 2008 Apr;52(4):1221–9. doi: 10.1128/AAC.01164-07. Epub 2008 Jan 22. PMID: 18212094; PMCID: PMC2292509.

51. Boakes E, Kearns AM, Ganner M, Perry C, Warner M, Hill RL, Ellington MJ. Molecular diversity within clonal complex 22 methicillin-resistant Staphylococcus aureus encoding Panton-Valentine leukocidin in England and Wales. Clin Microbiol Infect. 2011 Feb;17(2):140–5. doi: 10.1111/j.1469-0691.2010.03199.x. PMID: 20167006.

52. Coombs GW, Baines SL, Howden BP, Swenson KM, O’Brien FG. Diversity of bacteriophages encoding Panton-Valentine leukocidin in temporally and geographically related Staphylococcus aureus. PLoS One. 2020 Feb 10;15(2):e0228676. doi: 10.1371/journal.pone.0228676. PMID: 32040487; PMCID: PMC7010278.

53. Sánchez-Osuna M, Cortés P, Llagostera M, Barbé J, Erill I. Exploration into the origins and mobilization of dihydrofolate reductase genes and the emergence of clinical resistance to trimethoprim. Microb Genom. 2020 Nov;6(11):mgen000440. doi: 10.1099/mgen.0.000440. PMID: 32969787; PMCID: PMC7725336.

54. Gómez-Sanz E, Haro-Moreno JM, Jensen SO, Roda-García JJ, López-Pérez M. The Resistome and Mobilome of Multidrug-Resistant Staphylococcus sciuri C2865 Unveil a Transferable Trimethoprim Resistance Gene, Designated dfrE, Spread Unnoticed. mSystems. 2021 Aug 31;6(4):e0051121. doi: 10.1128/mSystems.00511-21. Epub 2021 Aug 10. PMID: 34374564; PMCID: PMC8407400.

55. Bukowski M, Banasik M, Chlebicka K, Bednarczyk K, Bonar E, Sokołowska D, Żądło T, Dubin G, Władyka B. Analysis of co-occurrence of type II toxin-antitoxin systems and antibiotic resistance determinants in Staphylococcus aureus. mSystems. 2025 Mar 18;10(3):e0095724. doi: 10.1128/msystems.00957-24. Epub 2025 Feb 27. PMID: 40013794; PMCID: PMC11915791.

56. Ramsay JP, Parahitiyawa N, Mowlaboccus S, Mullally CA, Yee NWT, Shoby P, Colombi E, Tan HL, Pearson JC, Coombs GW. Genomic characterization of a unique Panton-Valentine leucocidin-positive community-associated methicillin-resistant Staphylococcus aureus lineage increasingly impacting on Australian Indigenous communities. Microb Genom. 2023 Dec;9(12):001172. doi: 10.1099/mgen.0.001172. PMID: 38117559; PMCID: PMC10763498.

57. Al-Trad EI, Chew CH, Che Hamzah AM, Suhaili Z, Rahman NIA, Ismail S, Puah SM, Chua KH, Kwong SM, Yeo CC. The Plasmidomic Landscape of Clinical Methicillin-Resistant Staphylococcus aureus Isolates from Malaysia. Antibiotics (Basel). 2023 Apr 9;12(4):733. doi: 10.3390/antibiotics12040733. PMID: 37107095; PMCID: PMC10135026.

58. Dorado-Morales P, Garcillán-Barcia MP, Lasa I, Solano C. 2021. Fitness Cost Evolution of Natural Plasmids of Staphylococcus aureus. mBio 12:10.1128/mbio.03094-20. 10.1128/mbio.03094-20

59. Xue H, Wu Z, Li L, Li F, Wang Y, Zhao X. 2015. Coexistence of heavy metal and antibiotic resistance within a novel composite staphylococcal cassette chromosome in a Staphylococcus haemolyticus isolate from bovine mastitis milk. Antimicrob Agents Chemother 59:5788–5792. doi:10.1128/AAC.04831-14.

60. Harmer CJ, Hall RM. IS26 Family Members IS257 and IS1216 Also Form Cointegrates by Copy-In and Targeted Conservative Routes. mSphere. 2020 Jan 8;5(1):e00811–19. doi: 10.1128/mSphere.00811-19. PMID: 31915227; PMCID: PMC6952201.

61. Anne-Merethe Hanssen, Johanna U. Ericson Sollid, SCCmec in staphylococci: genes on the move, FEMS Immunology & Medical Microbiology, Volume 46, Issue 1, February 2006, Pages 8–20, 10.1111/j.1574-695X.2005.00009.x

62. Alcock, et al. 2023. CARD 2023: Expanded Curation, Support for Machine Learning, and Resistome Prediction at the Comprehensive Antibiotic Resistance Database. Nucleic Acids Research, 51, D690–D699.

